# LPCAT2 Regulates CD14 Expression During Macrophage Inflammatory Response to E. coli O111:B4

**DOI:** 10.1101/2023.06.27.546434

**Authors:** Victory Ibigo Poloamina, Wondwossen Abate, Gyorgy Fejer, Simon K Jackson

## Abstract

LPCAT2 is a lipid-modifying enzyme that co-localises in lipid rafts with TLR4 and regulates macrophage inflammatory response; however, its effect on TLR4 co-receptor–CD14 is unknown. RAW264.7 cells, a common murine macrophage experimental model, were used to study the effect of LPCAT2 on CD14 expression. The expression of LPCAT2 in RAW264.7 cells was silenced using RNA interference and treated with 100ng/ml of various lipopolysaccharide chemotypes. We found that CD14 expression induced by smooth lipopolysaccharide was significantly decreased (p < 0.05) in RAW264.7 macrophages with LPCAT2 silenced. This study suggests that LPCAT2 regulates CD14 gene and protein expression. This implies that LPCAT2 can regulate CD14-dependent cellular activities.

## Introduction

Inflammation is central to diseases such as cancer, asthma, sepsis, and cardiovascular disease and can be induced by stimuli such as cell injury, toxins, and pathogens [49]. Innate immunity which defines the first line of defense against microbial infection is characterised by the influx of inflammatory cells such as neutrophils, natural killer cells, and macrophages. Then, pattern recognition receptors like TLR4 (Toll Like Receptor 4) found on innate immune cells interact with pathogen associated molecular patterns like bacterial lipopolysaccharide to induce downstream processes that will result in the release of inflammatory cytokines [51].

Bacterial lipopolysaccharides (LPS) exist in various forms or chemotypes -the complete molecule that includes the ‘O-antigen ‘polysaccharide repeat units is referred to smooth (S) LPS, whereas various truncated forms are known as rough (R) form LPS and have subtypes referred to as Ra, Rc, Rd, and Re depending on the structure of the ‘core ‘region of the molecule [8-11].

CD14 is established as a co-receptor to TLR4 and MD2 for the recognition of bacterial lipopolysaccharide [7], it can be found anchored to macrophage membranes by a glycosylphosphatidylinositol [1 -3] or shed from cell membranes as soluble CD14 [4,5]. Both forms of CD14 are important to innate immune responses to infection in macrophages. Macrophages are primary effector cells of the innate immune system and secrete cytokines that attract other immune cells and initiate inflammation in response to pathogens at the site of infection [6]. CD14 is pivotal to endocytosis [12], MyD88-dependent and TRIF-dependent signalling [13,14], and phosphatidylinositol 4,5-biphosphate (PIP2) generation in lipid rafts [15,16].

LPCAT2 (Lysophosphatidylcholine Acyltransferase), also known as LysoPAF acetyltransferase localises in endoplasmic reticulum, lipid rafts, and lipid droplets in cell membranes [17,18]. It is physiologically relevant for asymmetry and fluidity of phospholipid membranes of inflammatory cells because it is part of the phospholipid remodeling pathway [50]. LPCAT2 is expressed on inflammatory cells such as peritoneal macrophages, microglia, and neutrophils [18, 43], it regulates inflammation by synthesising fatty acids and lipid mediators such as PAF, Leukotriene B4, and Prostaglandin E2 [19]. Although LPCAT2 is a lipid-modifying enzyme, it has also been shown to regulate the expression and modification of proteins involved in macrophage inflammatory responses to lipopolysaccharide [18, 20]. Several publications have suggested LPCAT2 as a biomarker for inflammatory disorders such as sepsis and allergic asthma [40 -42]. Furthermore, in experimental models of inflammatory diseases such as peripheral nerve injury and experimental allergic encephalomyelitis, LPCAT2 was significantly expressed [43 -47]. Experimental models deficient in LPCAT2 exhibit resolution of inflammatory responses and attenuation of neuropathic pain [18, 43, 48].

Despite evidence that LPCAT2 influences macrophage inflammatory response, the mechanisms of actions are not fully understood. This study shows that LPCAT2 can regulate CD14 expression, which will influence inflammatory processes that depend on CD14.

## Results

Endocytic process of bacterial LPS occurs between 0 – 1h [38, 39]. Therefore, the expression of CD14 in this paper has been analyzed at 30 minutes to correspond with the time course of endocytosis after infecting RAW264.7 cells with bacterial LPS. We have previously published the efficiency of LPCAT2 knockdown and shown that the cell manipulation does not influence housekeeping genes [20].

### 1. LPCAT2 gene silencing does not influence CD14 cell surface expression

We quantified the cell surface expression of CD14 at various time points after the LPS treatment of RAW264.7 cells. Figure 1 shows that LPS increases CD14 cell surface expression at 30 minutes and 24 hours (p≤0.0002). Moreover, there is a significant down-regulation in CD14 cell surface expression after 30 minutes (p≤0.04) and a gradual increase towards 24 hours (Figure 1A&C). In figure 1B, there is no significant difference between negative controls (p = 0.13); however, treating the cells with LPS caused a significant increase in CD14 cell surface expression (p=0.04). In figure 1D, there was no significant difference in LPS-induced CD14 cell surface expression after silencing LPCAT2 (p = 0.48).

**Figure 1:**
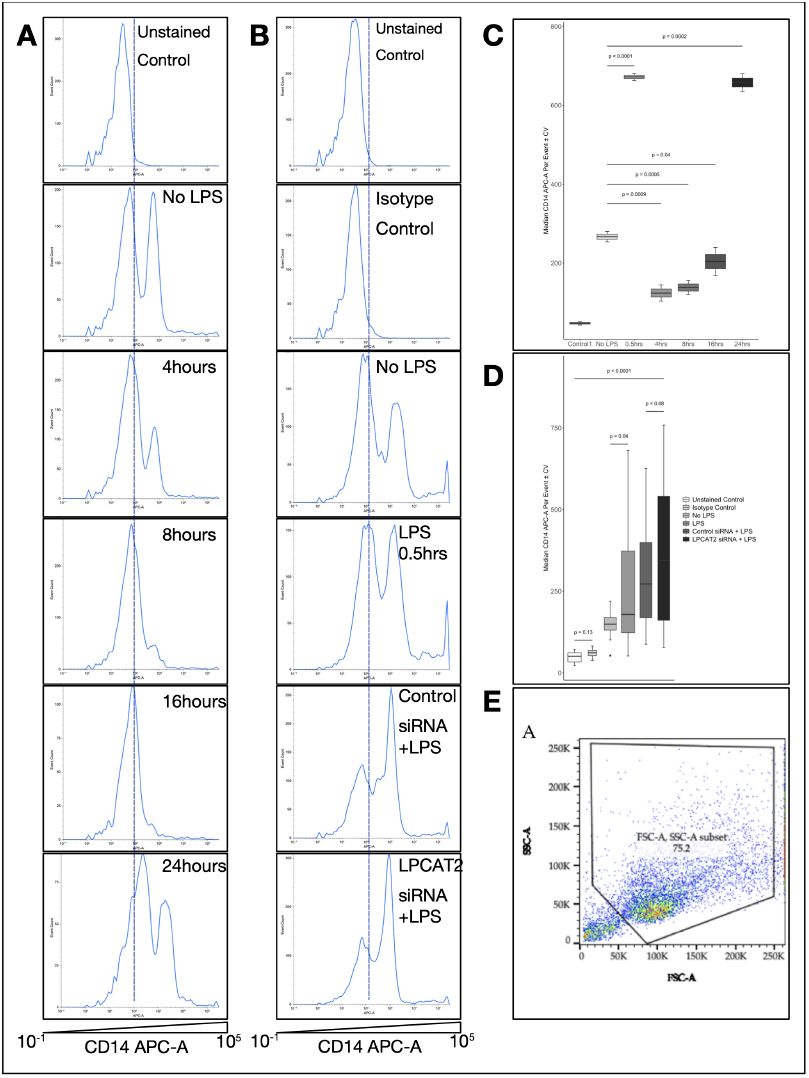
LPCAT2 Gene Silencing Does Not Influence Smooth LPS-induced CD14 Cell Surface Expression in RAW264.7 cells. RAW264.7 cells were transfected with 7nM LPCAT2 siRNA and treated with 100ng/ml smooth LPS (E. coli O111:B4) for 30 minutes to 24hrs. CD14 cell surface expression was determined with APC-labelled anti-mouse CD14 antibodies. **A –** Time course of LPS-induced CD14 cell surface expression in RAW264.7 cells. **B –** Effect of LPCAT2 silencing on LPS-induced CD14 cell surface expression. **C –** Box plots showing the time course of CD14 cell surface expression in LPS-treated RAW264.7 cells. **D –** Box plots of showing the effect of LPCAT2 silencing on LPS-induced CD14 cell surface expression. **E –** Representative density plot of RAW264.7 cells. Data obtained from at least three independent experiments and represent Median ± CV of APC fluorescence area per event. Unstained control and Isotype control were negative controls. Control 1 is unstained control.

### 2. LPCAT2 gene silencing influences CD14 mRNA and protein expression

The mRNA expression of LPS receptor complex; CD14 and TLR4 was quantified before and after silencing LPCAT2 in RAW264.7 cells. LPCAT2 gene silencing did not influence TLR4 protein expression or mRNA expression, but TLR4 mRNA was significantly decreased in RAW264.7 cells treated with smooth LPS. On the other hand, like the cell surface expression, LPS significantly increased CD14 protein expression at 30 minutes (2.0±0.2, p=0.001). However, LPCAT2 gene silencing caused a significant decrease at 30 minutes (0.7±0.1, p = 0.01).

### 3. LPCAT2 gene silencing only influences O111:B4 LPS-induced mRNA expression of IL6, IP10, and IFNβ

The effect of LPCAT2 gene silencing on LPS-induced mRNA expression of IL6, IP10, and IFNβ was analyzed. All chemotypes of LPS increased the mRNA expression of IL6, IP10, and IFNβ. LPCAT2 gene silencing had no significant effect on the mRNA expression of the cytokines in untreated cells (Fig. 3A). However, LPCAT2 silencing significantly decreased the cytokine mRNA expression in cells treated with S-form LPS (Fig 3B) but not in cells treated with either R-form LPS (Fig 3C, D).

### 4. LPCAT2 gene silencing influences IL6, IP10, and IFNβ secretion

We measured the effect of silencing LPCAT2 on the amounts of secreted IL6, IP10, and IFNβ in cells treated with O111:B4 LPS. S-form LPS treatment significantly increased the secreted cytokine protein from the cells (Fig. 4). Moreover, Figure 4 shows that LPCAT2 silencing significantly decreased the secreted proteins in cells stimulated with S-form LPS.

## Discussion

Several publications have shown that LPCAT2 affects LPS-induced inflammatory responses in macrophages, and LPS increases either LPCAT2 expression or enzymatic activity [18,20,23]. The mechanism for LPCAT2 is yet to be fully understood. It is already well-established that CD14 mediates LPS-induced inflammatory responses in macrophages [24]. Since LPCAT2 affects LPS-induced inflammation in macrophages, we analysed the effect of LPCAT2 on the expression of LPS receptor complex proteins TLR4 and CD14. We found that silencing LPCAT2 expression in RAW264.7 cells resulted in reduced expression of CD14 mRNA and CD14 total cellular protein. On the other hand, LPCAT2 had no significant effect on TLR4 mRNA and protein expression (figure 2). Previous research suggested that TLR4 and CD14 expression were not affected by LPCAT2 [18]. This suggestion holds for TLR4 but not for CD14 in this study. However, as the publication did not show data, the study may be referring to the surface expression of CD14. As seen in figure 1, silencing LPCAT2 had no significant effect on CD14 surface expression. Unpublished work has previously shown that over-expression of LPCAT2 increases CD14 expression [25]; this supports the results in figure 2. An old study showed differential levels of CD14 expression, with smooth LPS being more potent at increasing CD14 expression than rough LPS [26]. Nonetheless, here, the difference is not seen with the potency of each LPS chemotype but in the effect of silencing LPCAT2 (figure 2A), this difference may be because of different experimental models used. The results show that silencing LPCAT2 much decreases smooth LPS-induced CD14, IL6, IP10, and IFNβ mRNA expression, but not rough LPS (figure 2A, figure 3). Table 1 shows that the percentage change in CD14 and cytokine mRNA expression decreases with increasing LPS roughness. LPCAT2 did not influence TLR4 mRNA expression; there is an apparent difference between the effect of smooth LPS and rough LPS on TLR4 mRNA expression. Smooth LPS-induced decrease of TLR4 mRNA expression is in line with previous studies [27, 28]. It will be interesting to know why rough LPS did not decrease TLR4 mRNA in future. Although LPCAT2 does not influence TLR4 protein expression, it influences its subcellular localisation and possibly its post-translational modification [18, 20]. Various other cellular mechanisms besides genetic transcription can determine the amount of secreted cytokines [29, 30]; therefore, the mRNA expression is comparably a more specific way to study the effects of the various chemotypes of LPS on cytokine expression. Despite that, silencing LPCAT2 had a similar effect on smooth LPS-induced cytokine secretion as its mRNA expression (figure 4).

**Figure 2:**
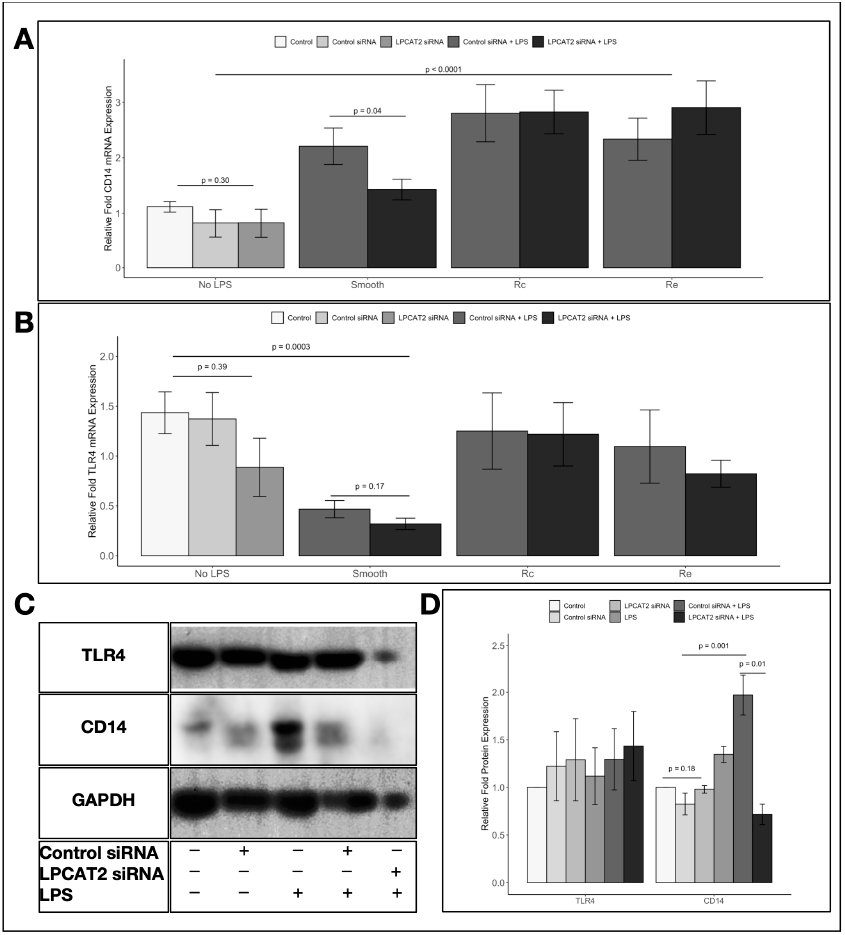
LPCAT2 gene silencing influences smooth LPS induced CD14 mRNA and protein expression in RAW264.7 Cells, but not TLR4 expression. RAW264.7 cells were transfected with 7nM LPCAT2 siRNA and treated with 100ng/ml smooth LPS (E. coli O111:B4), Rc LPS (E. coli J5), and Re LPS (E. coli K12 D32m4) for 30 minutes. mRNA expression was determined with RT-qPCR, and protein expression by western blot. **A –** CD14 mRNA expression. **B –** TLR4 mRNA expression. **C –** Western blots of TLR4 and CD14 proteins at 30 minutes. **D –** Western blots of CD14 protein at 30 minutes. Data from at least three independent experiments; bar and error bars show mean ±standard error of mean; + mean present, – means absent.

**Figure 3:**
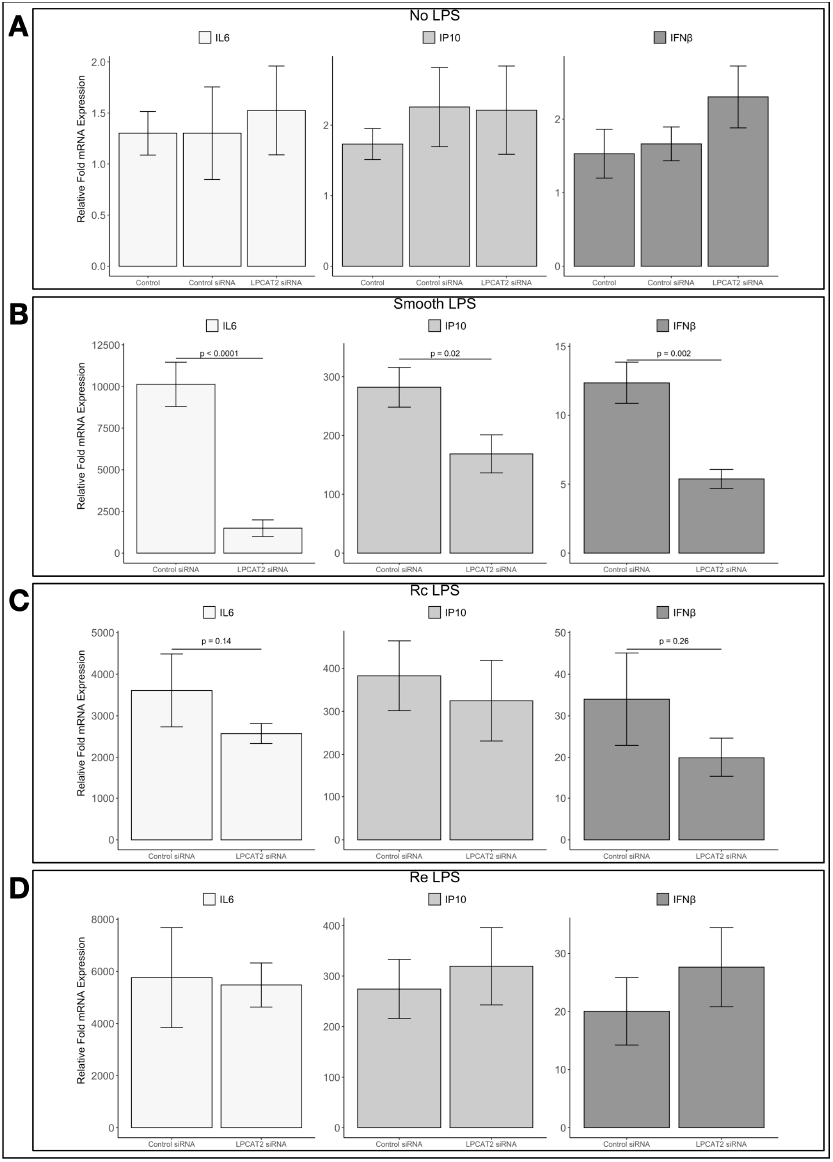
LPCAT2 gene silencing only influences O111:B4 LPS-induced mRNA expression of IL6, IP10, and IFNβ. RAW264.7 cells were transfected with 7nM LPCAT2 siRNA and treated with 100ng/ml smooth LPS (E. coli O111:B4), Rc LPS (E. coli J5), and Re LPS (E. coli K12 D32m4) for 6 hours. mRNA expression was determined with RT-qPCR. **A –** No ligand (or no LPS). **B –** O111:B4 LPS. **C –** J5 LPS. **D –** K12, D31m4 LPS. Data from at least three independent experiments; bar and error bars show mean ±standard error of mean.

**Figure 4:**
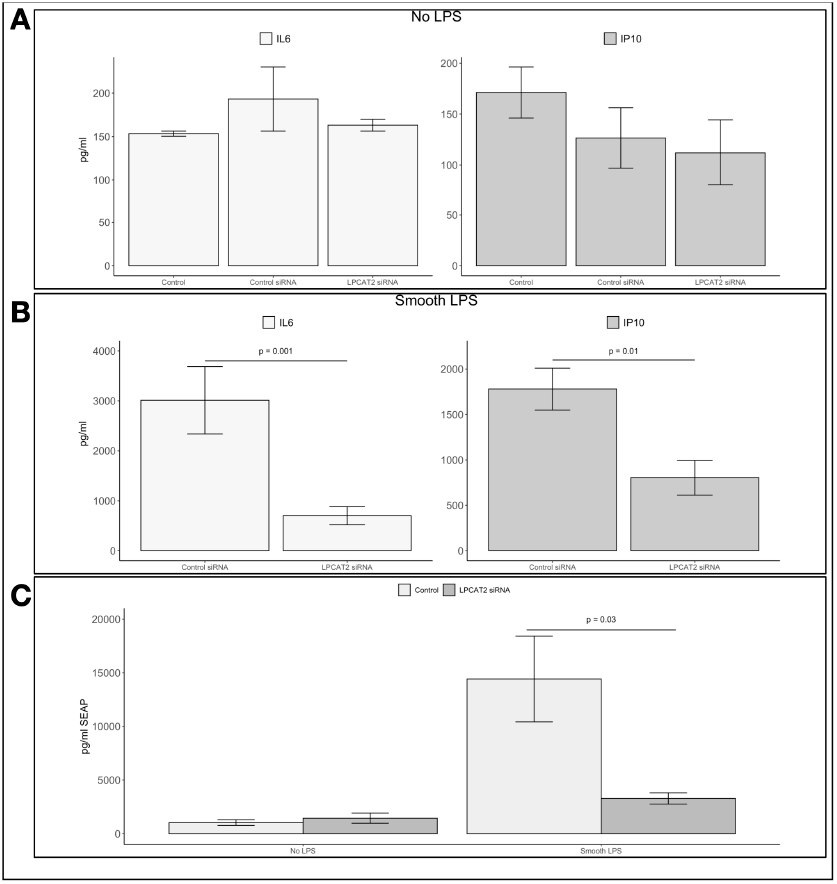
LPCAT2 gene silencing influences IL6, IP10, and IFNβ secretion: RAW264.7 cells were transfected with 7nM LPCAT2 siRNA and treated with 100ng/ml smooth LPS (E. coli O111:B4) for 24 hours (A&B) or 18hours (C). Cytokine secretion was measured with ELISA and RAW-Blue ISG reporter gene assay. **A –**No ligand (or no LPS). **B –** IL6 and IP10 secretion. **C –** IFNβ secretion. Data from at least three independent experiments; bar and error bars show mean ±standard error of mean.

**Figure 5:**
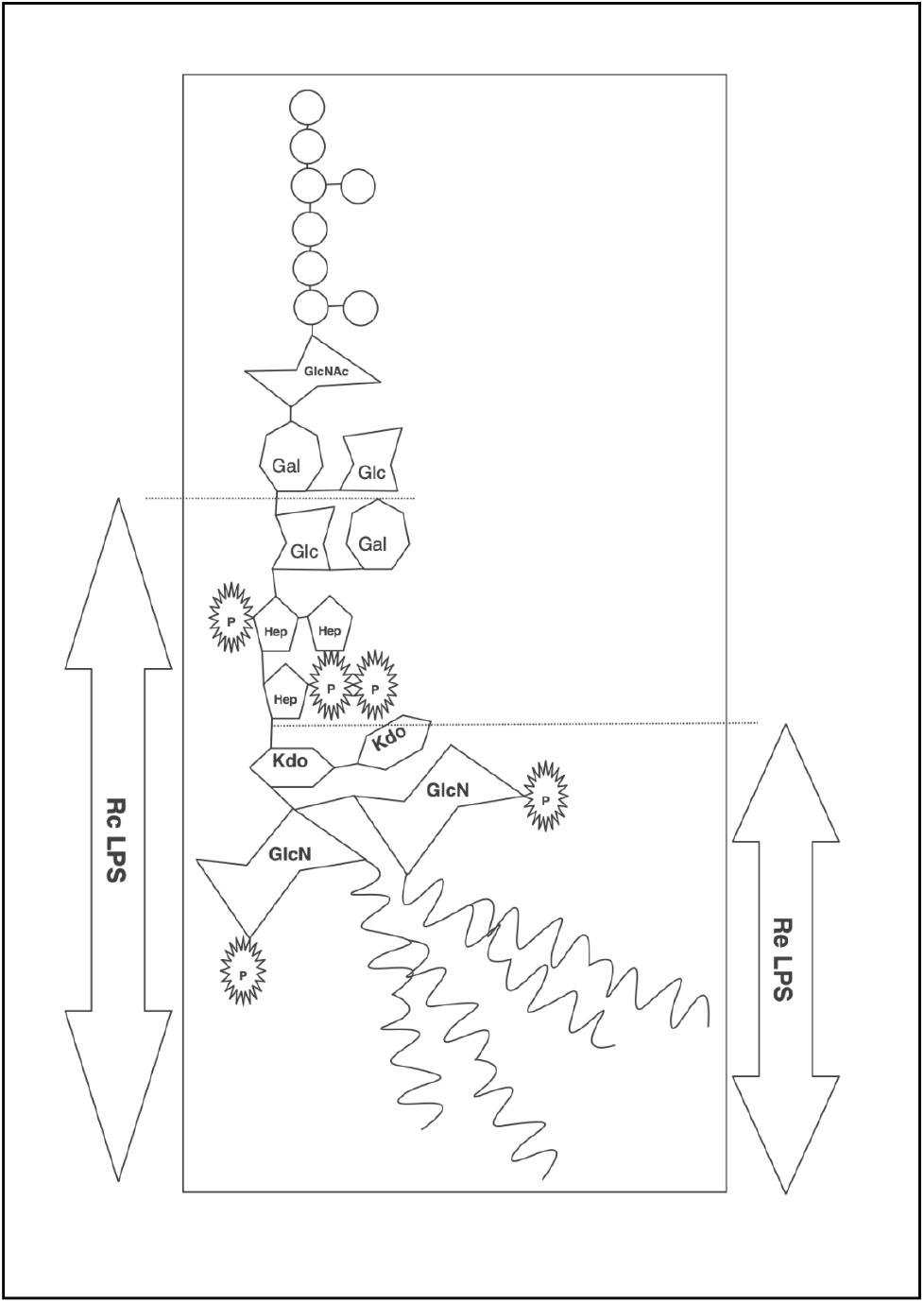
General Structure of Bacterial Lipopolysaccharide Showing Rc and Re Mutant Structures. p – phosphate group; GlcN – glucosamine; Kdo – 3-deoxy-D-manno-octulosonic acid; Hep – L-glycerol-D-manno-heptose; Glc – glucose; Gal – galactose; GlcNAc – N-acetyl-glucosamine. Circles represent the O-antigen region of LPS [8-11].

**Table 1:**
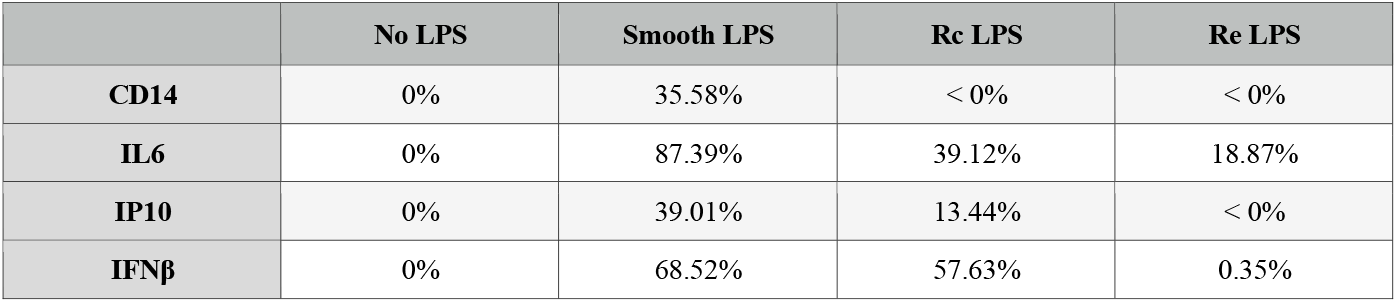
Percentage Decrease in CD14, IL6, IP10, and IFNβ mRNA Expression Per LPS Chemotype. Data from at least three independent experiments; Values show percentage decrease in mRNA expression after silencing LPCAT2 in RAW264.7 cells; calculated from fold change values in figure 3 and figure 2A. <0% indicates increased expression.

|  | No LPS | Smooth LPS | Rc LPS | Re LPS |
| --- | --- | --- | --- | --- |
| **CD14** | 0% | 35.58% | < 0% | < 0% |
| **IL6** | 0% | 87.39% | 39.12% | 18.87% |
| **IP10** | 0% | 39.01% | 13.44% | < 0% |
| **IFNβ** | 0% | 68.52% | 57.63% | 0.35% |

Both TRIF-dependent and MyD88-dependent signalling requires CD14 in macrophages. At low concentrations, rough LPS induces MyD88-dependent signalling independently of CD14. Whereas smooth LPS, considered a weaker activator, requires CD14 to induce inflammation [31-34]. CD36 can substitute for CD14 in loading rough LPS but not smooth LPS onto TLR4/MD2 signalling complex, allowing for CD14-independent macrophage responses to rough LPS [35]. The ability of LPCAT2 to regulate the expression of CD14, IL6, IP10, and IFNβ induced by 100ng/ml of smooth LPS but not rough LPS suggests that LPCAT2 regulates LPS-induced inflammation in a CD14-dependent manner. The ability of LPCAT2 to regulate total CD14 expression implies that LPCAT2 influences processes that depend on CD14. As aforementioned, one of these processes is smooth LPS-induced inflammation. Endocytosis, which occurs after LPS recognition, depends on CD14 [12]. Therefore, this study implies that LPCAT2 may influence endocytosis. Again, the clustering of CD14 in lipid raft compartments of the cell membrane triggers the generation of PIP2 [15]. PIP2 is a phospholipid that regulates MyD88-dependent TLR4 signalling, and its hydrolysis or phosphorylation enables TRIF-dependent TLR4 signalling [36, 37]. A previous study showed that silencing LPCAT2 reduced TLR4 expression in the lipid rafts [18]; therefore, it is possible that silencing LPCAT2 may regulate the clustering of the LPS receptor complex induced by LPS binding and attenuate TLR4 signalling.

The results from this study are compelling, but will the effects of LPCAT2 on CD14 expression and the differential response to LPS chemotypes be the same in human cells? To conclude, LPCAT2 regulates CD14 expression and cellular processes that depend on CD14 in RAW264.7 cells.

## Materials and Methods

Chemical reagents for preparing buffers and the BCA Assay Kit were purchased from Sigma Aldrich, UK and Fisher Scientific, UK. Buffers used include, phosphate-buffered saline (PBS), ELISA blocking buffer (0.1% bovine serum albumin in PBS-0.1% Tween 20), Western transfer blocking buffer (5% bovine serum albumin in Tris Buffered Saline (TBS)), cell lysis buffer (TBS-1% Triton X-100), SDS sample buffer (187.5mM Tris-HCl -pH 6.8, 6% SDS, 30% glycerol, 0.03% bromophenol blue, 100mM beta-mercaptoethanol), and enhanced chemiluminescence detection reagent (20mM Tris-HCl-pH 7.5, 250mM Luminol, 90mM p-Coumaric acid, 1:2500 Hydrogen Peroxide). PolyPlus INTERFERin was purchased from Source Bioscience, UK. Pre-designed siRNA, Opti-MEM, Power SYBR Green, RNA to cDNA kit, and gel casting materials were purchased from Life Technologies, UK. Antibodies for western blot and flow cytometry, Protein A/G agarose gel beads, and protein ladders were obtained from Santa Cruz Biotechnology, UK [sc-516646, sc-293072, sc-516102, sc-32233, sc-2020, sc-358914], R&D Technologies [1574893S, 16789972] and BD Pharminogen[560634]. DMEM culture medium and other cell culture materials were purchased from Lonza, UK. PCR primers were designed with Primer3 Plus Bioinformatics Software and NCBI BLAST and purchased from Eurofins Genomics. Recombinant mouse proteins, ELISA antibodies and detection reagents were purchased from EBioscience [14-7061-81], and Peprotech Ltd. [500-P129]. Smooth LPS – E. coli O111:B4, Sigma-Aldrich L260; Rc LPS – E. coli J5, List Biological Laboratories, 301; Re LPS – E. coli K12 D32m4, List Biological Laboratories, 302]. LAL reagent water [Lonza, W50-640]. BCA Assay kit [ThermoFisher Scientific, 23227].

### 1. Cell line and Culture

RAW264.7 cell line was obtained from the European Collection of Cell Cultures (ECACC) through Public Health England, UK. RAW264.7 macrophages were maintained in Dulbecco’s Modified Eagle Medium (DMEM) [Lonza, BE12-914F] supplemented with 10%(v/v) Foetal Bovine Serum (FBS) [Labtech.com, BS-110] and 1%(v/v) 0.2M L-Glutamine [BE17-605E], and incubated at 37ºC, 5% CO_2_. RAW-Blue ISG and Quantified-Blue assay kit were obtained from Invivogen, Ltd. The cell culture medium for RAW-Blue ISG contained 2%(v/v) Normocin and Zeocin.

### 2. Preparation of Lipopolysaccharide

Lipopolysaccharide was resuspended in LAL reagent water (<0.005EU/ml endotoxin levels). Cells were stimulated with 100ng/ml lipopolysaccharide in cell culture medium.

### 3. Transfection of RAW264.7 Cells with LPCAT2 siRNA and RNF19B siRNA

RAW264.7 cells were cultured 24 hours before gene silencing. Then using Opti-MEM (Reduced Serum Medium) as a diluent, a transfection mixture containing 7nM of siRNA was prepared and added to cells. Finally, the cells were incubated at 37ºC, 5% CO_2_ with Opti-MEM for 24 hours for efficient gene silencing.

### 4. Sandwich ELISA for Quantifying Secreted IL6 and IP10 Proteins

The medium from RAW264.7 cells stimulated with lipopolysaccharide was analysed for the quantity of secreted IL6 and IP10 after 24 hours. ELISA was carried out according to manufacturer’s instructions.

### 5. Reverse Transcription and Real-Time Quantitative PCR

Gene expression of cytokines was analysed after 6 hours of treatment with 100ng/ml LPS. The reaction master mix contained 37%(v/v) nuclease-free, 230nM of target primers (a mixture of both forward and reverse primers), 60%(v/v) Power SYBR Green, and 3μl of ≥ 100ng/μl cDNA. The reaction was initiated at 95ºC for 10 minutes, then up to 40 repeated cycles of denaturing (15 seconds, 95ºC), annealing, and extension (60 seconds, 60ºC). GAPDH and ATP5B were used as endogenous control.

**Table 2:**
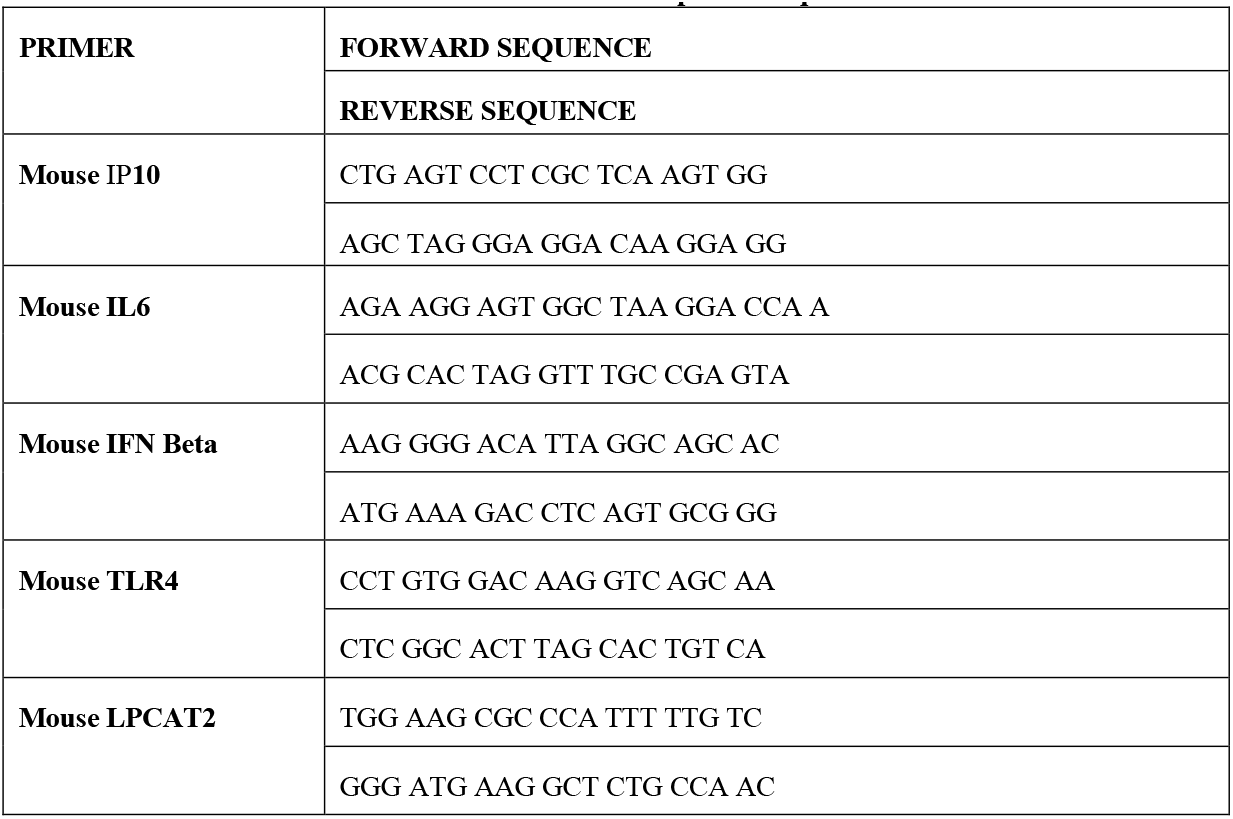
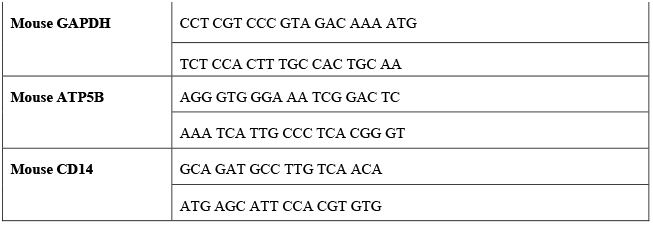
List of primer sequences.

### 6. Western Blot Analysis

Equal amounts of whole-cell lysates were separated on pre-cast SDS-PAGE gels and blotted onto a PVDF membrane using a blot module. The blots were blocked with blocking buffer, probed with primary and HRP-conjugated secondary antibodies. The target proteins separated on the PVDF membrane were detected and analysed using ImageJ.

### 7. RAW-Blue™ ISG Reporter Gene Assay

RAW-Blue ISG cell lines are derived from RAW264.7 cell line, designed to stably express secreted embryonic alkaline phosphatase (SEAP) gene which is controlled by interferon stimulated gene 54 (ISG54)-inducible promoter. Stimulation of the cell with type I IFN inducers causes productions of SEAP, which is quantified with QUANTI-Blue™ [21].

To quantify type I IFN after stimulating RAW264.7 cells with 100ng/ml lipopolysaccharide, RAW-Blue ISG cells were incubated with medium from stimulated RAW264.7 cells for 18 hours. Medium containing recombinant mouse IFN beta was used as standards for this assay. After 18 hours, SEAP secreted by RAW-Blue ISG cells into the medium was quantified with Quanti-Blue according to manufacturer’s instructions [Invivogen].

### 8. Flow Cytometry Analysis

100000 Live RAW264.7 cells were rinsed with and resuspended in 4ºC PBS. They were counted using trypan blue exclusion method. The live cells were incubated at 4ºC with target antibodies for 30 minutes and resuspended in 4ºC PBS to wash off unbound antibodies. Dead cells and unbound antibodies were excluded by centrifugation (200xg 5minutes). The antibody-bound cells were then resuspended in 4ºC 5%(v/v) FBS in PBS and used for flow cytometry. A 0.7micron insert was used on BD FACSAria II flow cytometer. Data was analysed and visualised using FACSDiva software [BD, UK] and Floreada.io web-based software.

### 9. Data and Statistical Analysis

Statistical analysis was carried out in graphpad prism. Independent experiments were repeated at least three times. Data represent Mean ±Standard Error of Mean unless stated otherwise. Paired T-Test with Dunnett’s T3 multiple comparison tests was used for statistical analysis. All statistical tests were significant at a 95% confidence interval, p ≤ 0.05. Relative quantification of mRNA expression was carried out using the 2^-ΔΔCt^ method [22], GAPDH and ATP5B were used as endogenous reference genes.

## Declarations

### Ethics approval and consent to participate

Not Applicable

## Availability of Data and Material

Supplementary data have been submitted with this manuscript. For more information on data availability, please contact the corresponding author at the email address provided.

## Author Contributions

Prof. Simon K. Jackson, Dr Wondwossen Abate, and Dr Gyorgy Fejer reviewed and supervised this research. The research was carried out and written up by Dr Victory Ibigo Poloamina.

