## supplementary data for "LPCAT2 Regulates CD14 Expression During Macrophage Inflammatory Response to E. coli O111:B4"

### 1. TLR4 Expression after silencing LPCAT2 in RAW264.7 cells

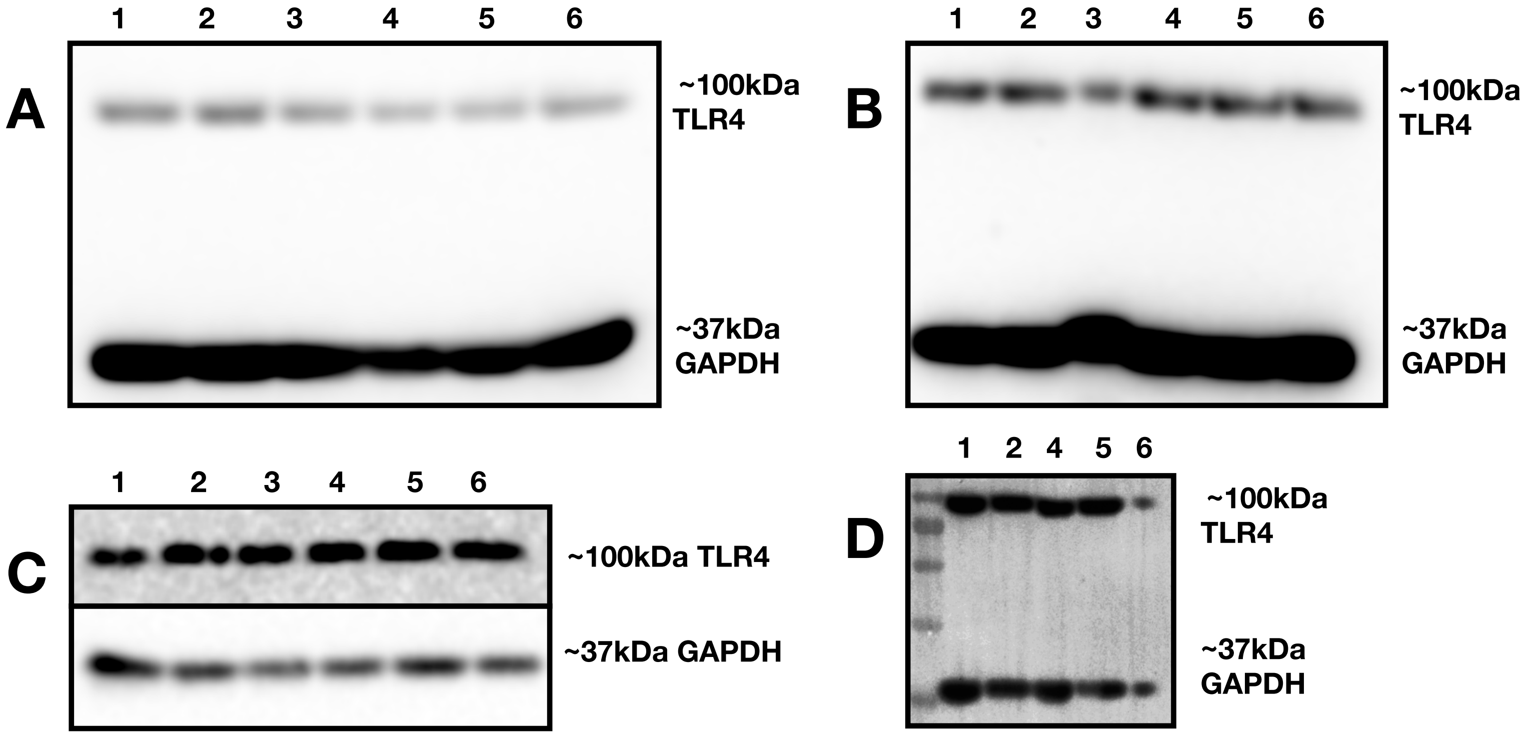


**Supplementary Figure 1: TLR4 Expression before and after silencing LPCAT2 and treatment with 100ng/ml LPS for 30 minutes.** Lane 1- Untreated (Medium only), Lane 2- Negative siRNA, Lane 3 - LPCAT2 siRNA, Lane 4- LPS, Lane 5- Negative siRNA + LPS, Lane 6- LPCAT2 siRNA + LPS. Each subfigure represents an independent experiment.

#### 2. CD14 Expression after silencing LPCAT2 in RAW264.7 cells

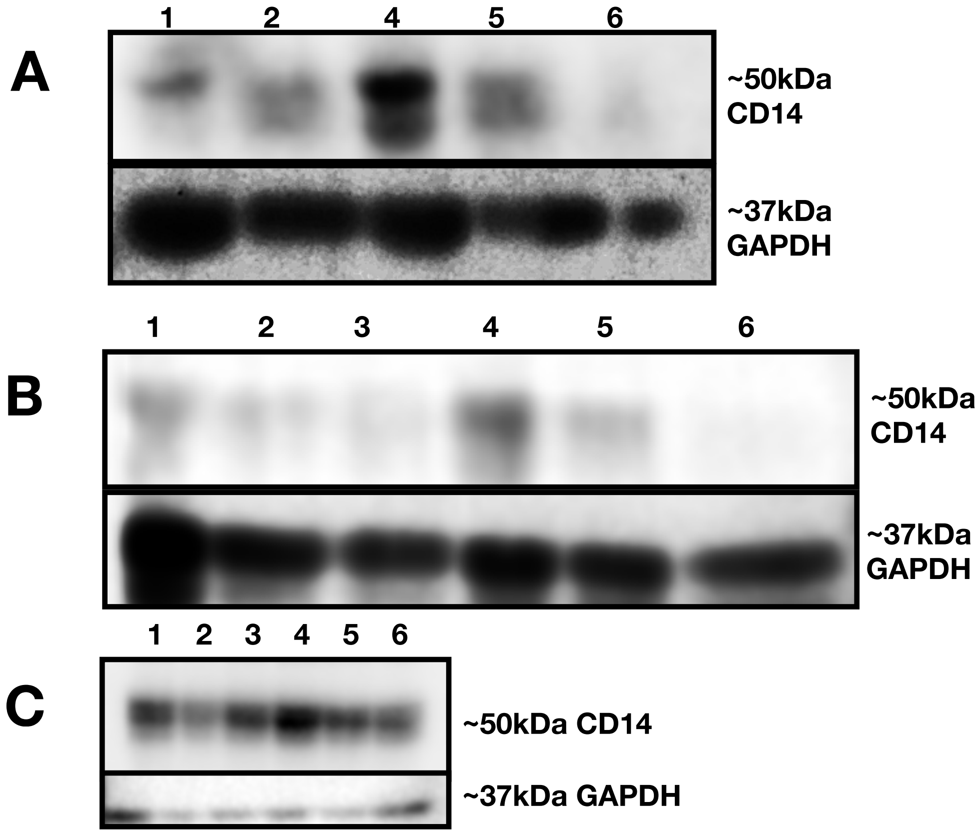


**Supplementary Figure 2: CD14 Expression before and after silencing LPCAT2 and treatment with 100ng/ml LPS for 30 minutes.** Lane 1- Untreated (Medium only), Lane 2- Negative siRNA, Lane 3 - LPCAT2 siRNA, Lane 4- LPS, Lane 5- Negative siRNA + LPS, Lane 6- LPCAT2 siRNA + LPS. Each subfigure represents an independent experiment.
